# Implicit and explicit learning of Bayesian priors differently impacts bias during perceptual decision-making

**DOI:** 10.1101/2021.03.05.434141

**Authors:** V. N. Thakur, M. A. Basso, J. Ditterich, B. J. Knowlton

## Abstract

Knowledge without awareness, or implicit knowledge, influences a variety of behaviors. It is unknown however, whether implicit knowledge of statistical structure informs visual perceptual decisions or whether explicit knowledge of statistical probabilities is required. Here, we measured visual decision-making performance using a novel task in which humans reported the orientation of two differently colored translational Glass patterns; each color associated with different orientation probabilities. The task design allowed us to assess participants’ ability to learn and use a general orientation prior as well as a color specific feature prior. Classifying decision-makers based on a questionnaire revealed that both implicit and explicit learners implemented a general orientation bias by adjusting the starting point of evidence accumulation in the drift diffusion model framework. Explicit learners additionally adjusted the drift rate offset. When subjects implemented a stimulus specific bias, they did so by adjusting primarily the drift rate offset. We conclude that humans can learn priors implicitly for perceptual decision-making and depending on awareness implement the priors using different mechanisms.

---

Regularities in the environment can be learned implicitly and can bias judgments, preferences or fluency of movement^1–7^. For example, the statistical regularities of one’s native language are readily acquired by infants ^8^. Humans are also able to learn to use contextual information to search the location of a target in a display when the location is correlated with contextual features. Contextual cuing can occur without awareness of the correlation^9^. Implicit learning of regularities in finite-state artificial grammars also occurs through exposure to exemplars formed according to the grammar. After such exposure, participants classify new items as grammatical despite lacking awareness for the grammatical rules ^5,10–12^. Artificial grammar learning is intact in patients with amnesia, indicating independence from declarative learning. Perceptuo-motor behavior is also influenced by implicitly learned statistical regularity. In the Serial Reaction Time task, participants press keys on a keypad corresponding to cued locations. After practice, participants show reduced reaction times for locations occurring according to a sequence despite a lack of explicit knowledge of the sequence. Amnesic patients who have no awareness of the learning, show similar levels of sequence learning in this task compared to healthy control participants ^13,14^.

Although implicit knowledge can influence cognitive processes like attention-shifting and judgments of goodness-of-fit in an artificial grammar learning procedure, it is less clear whether implicit knowledge could become integrated with diagnostic information that is present to influence a judgement. In many circumstances, base rates, or Bayesian priors, influence judgements^15–23^. For example, a cloudy sky in Seattle might lead one to grab an umbrella when leaving home, while the same sky would not lead to this decision in Los Angeles. Here, we investigate whether base-rate priors can be learned implicitly and can influence perceptual judgements. On one hand, much of our knowledge of regularities of the environment is learned implicitly, and thus it seems adaptive that such knowledge should contribute to judgments. On the other hand, implicit knowledge of priors may not integrate readily with perceptual decisions that are based on sensory information in awareness. Here, we test the hypothesis that base rate priors that are implicitly acquired can influence orientation judgments.

If implicit knowledge of priors influences perceptual judgments, it may be that this knowledge affects judgments differently than explicit knowledge of priors. To compare decision-making under these different conditions, we applied the Drift Diffusion Model (DDM) to behavioral data and assessed how model parameters shift when subjects are aware of priors compared to when priors are implicitly learned.

## RESULTS

Participants made judgments about the orientation (45˚ left or right) of a dynamic Glass pattern stimulus^24^. We parameterized the difficulty of the perceptual judgement on each trial by varying the dot pair correlations, referred to as coherence, ranging from 0 - 100% with 0% having no dot pair correlations and thus no orientation signal and with 100% having all dot pairs correlated and thus the strongest orientation signal. On each trial, after the appearance of a centrally-positioned fixation spot for a fixation time of 1000±200ms, two choice targets appeared, one on the left and the other on the right of the screen, to orient the participants. Then, a Glass pattern stimulus appeared and the fixation spot disappeared and the participants reported their perception - left or right - with a key press; either ‘o’ for left or ‘p’ for right. If the participant chose correctly, an audible tone provided feedback. Incorrect choices received no tone and the 0% coherence trials received feedback consistent with the base-rate prior for that condition (e.g., 50% of the trials in the condition in which the two orientations were equally likely; Figure 1a).

**Figure 1:**
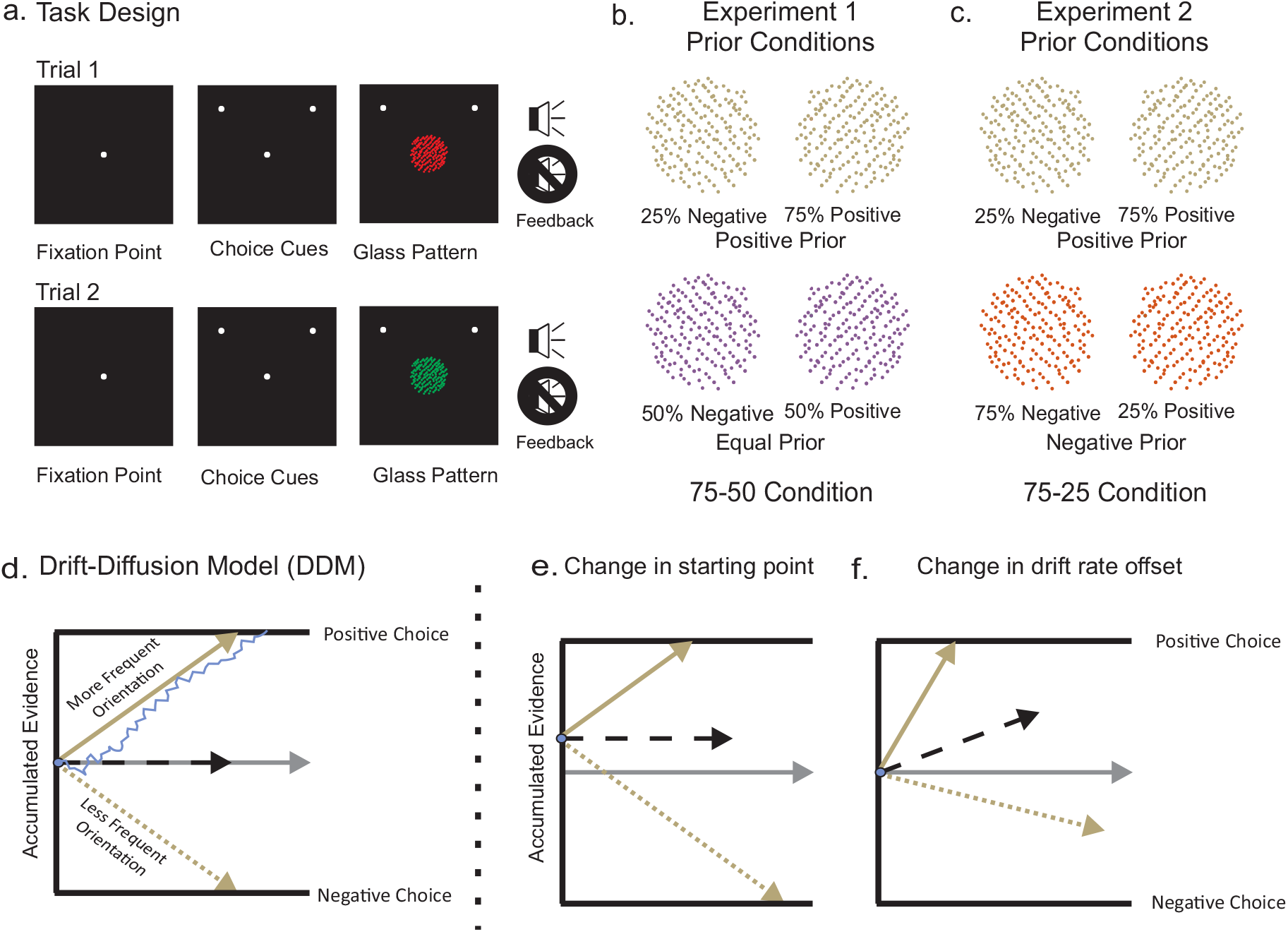
Schematics of the task design and the drift diffusion model (DDM). a. The black squares show the screen that the participants viewed and illustrate the spatial and temporal arrangements of the task. The white circle in the center of the screen shows the fixation spot and the two additional white circles indicate the two possible choices; left or right. The fixation spot and the choice targets appeared sequentially with a delay of 1000ms ± 200ms between. Next the Glass pattern appeared and the fixation spot disappeared simultaneously, cueing the participant to report the perceived orientation with a key press (‘o’ for left and ‘p’ for right). A tone, indicated by the audio symbol, provided feedback only for correct choices. b. Experiment 1. Khaki and lilac Glass patterns (100% coherence, 45 deg orientation negative and positive), illustrate the prior manipulation. The negative orientation occurred for the khaki Glass pattern on 25% of the trials and the positive orientation occurred on the remaining 75% of the khaki Glass pattern trials (Positive Prior). For the lilac Glass pattern stimuli, the negative and positive orientations occurred with equal probability (Equal Prior). All trials types were interleaved randomly and we counterbalanced the color and direction of the Glass pattern stimuli across participants and eventually converted to positive and negative orientation for analysis purpose. c: Experiment 2: The negative orientation occurred on 25% of the khaki Glass pattern trials and the positive orientation occurred on the remaining 75% (Positive Prior). In the same experimental block, the negative orientation occurred on 75% of the orange Glass pattern trials and the positive orientation occurred on remaining 25% of the orange Glass pattern trials (Negative Prior). All trial types were randomly interleaved and we counterbalanced the color and direction of the Glass pattern stimuli across participants and eventually converted into positive and negative orientations for analysis purpose. See text and methods for further details. d. The DDM of perceptual decision making proposes that sensory evidence is accumulated over time until a bound is reached, here labeled Positive and Negative. The solid khaki arrow shows the drift rate of evidence accumulation, labeled drift rate for more frequent orientations and the dashed khaki arrow shows the same for less frequent orientations. The blue line shows the noisy evidence accumulation for an individual trial. In the absence of a bias, evidence for either the positive or the negative decision begins to accumulate at a point that is equidistant from the two bounds, referred to as the starting point of evidence accumulation (khaki arrow). Time is indicated along the horizontal axis. e. Same as in d except the accumulation process begins at a point closer to Positive decision bound (shown by dashed black arrow). f. Same as in d except with an increased drift rate offset for positive decisions (shown by dashed black arrow)

The critical aspect of this task was that each Glass pattern stimulus appeared in one of two different colors (red or green) and for each participant, the probability of the left or right orientation for each color differed. Thus, stimulus color provided information that could contribute to the perceptual decision about orientation, and in this sense knowledge of the color-orientation relationship acts as a Bayesian prior ^25,26^. The stimulus color and prior probabilities were counterbalanced across participants, hence we converted the orientations to positive (right) and negative (left) orientations. In Experiment 1, referred to as the 75-50 condition, one colored Glass pattern had positive orientation priors and the other had equal orientation priors (Fig. 1b). Similar to Experiment 1, Experiment 2 also used two colored Glass patterns with different prior probabilities (Fig. 1c). In Experiment 2, each color was associated with a positive prior and negative prior with equal strength, so there was no base-rate difference for the two orientations across all stimuli. Previous studies on perceptual decision-making have not directly examined whether knowledge of priors was explicit or implicit. Here, we were able to classify participants’ knowledge of the priors based on their responses to a questionnaire given after the session, and compare their performance to that of participants who were informed of the priors at the onset of the task.

To characterize differences based on the type of knowledge of the priors, we examined performance using the DDM. The DDM considers a process where the net evidence – the difference in evidence for or against a particular outcome - accumulates with time. When the amount of evidence reaches a level associated with one or the other bound or threshold, a decision is made (Fig. 1d). In this framework, priors can create biases in decision-making either by changing the starting point from which evidence begins to accumulate to be closer to the bound associated with the more frequent alternative^17,19,27,28^ (Figure 1e), or by increasing the rate of evidence accumulation for the more frequent alternative ^1,23,29,30^ (Figure 1f), or both^21,26,31^. We modeled the data obtained from Experiments 1 and 2 using a simple DDM (see Methods) to assess the mechanisms underlying how priors create perceptual decision biases in implicit and explicit learners.

### Experiment 1 - Participants Learn Priors Implicitly and Explicitly

Data from 67 participants were analyzed in Experiment 1, all with informed consent using procedures approved by the UCLA IRB. Based on responses to the questionnaire, participants were divided into three groups. Participants were classified as “Implicit Learners ” if they reported no awareness that the different colors were associated with different orientation base-rates and that they perceived the orientations to occur with equal frequency (N=23, 34%). Participants were classified as “Partial Explicit Learners ” if they reported an overall difference in the orientation frequency but were unaware that this differed by color or could only identify the orientation prior for one color (N=20, 30%). A group of participants (N=24, 37%) classified as ‘Explicit Learners’ was informed of the priors for the different colored stimuli at the outset of the session and they all reported them correctly at the end of the session.

All participants performed the task well, using the orientation information to guide their choices (Fig. 2a-c). All participants also showed a bias in making more positive choices. We calculated the bias by measuring the proportion of choices made at 0% coherence. For the Implicit Learners, the mean proportion of positive choices on the positive prior trials was 0.575, (Figure 2a, difference from chance: t(22)=31.88, p<0.001). Furthermore, there was a 4.2% increase in positive choices compared to chance level even for the equal prior trials (Figure 2a, 0.542; difference from chance: t(22)=25.25, p<0.001). The bias did not differ significanlty between the two conditions (t(22)=1.008, p=0.289). The Partial Explicit Learners showed a similar trend in performance and the use of priors, showing a significant 12.2% increase from 50% in choosing the positive orientation in the positive prior trials (Figure 2b, mean proportion = 0.622; t(19) = 24.09, p< 0.001). For the equal prior trials, the Partially Explicit Learners also showed a 5.1% increase in choosing the positive orientation (Figure 2b, mean proportion=0.551; t(19)=20.25, p<0.001), which was lower than for the positive prior trials (t(19)=2.274, p<.05). Thus, implicit learning of both the general prior (Implicit group) and the color-specific priors (Partial Explicit group) occurred in the participants who were not informed of the base rate differences. The mean bias for the positive prior trials in the Explicit group was 0.645, which was significantly above chance (Figure 2c, t(23)=28.08, p<0.001). The Explicit Learners also showed 5.8% increase in bias for the equal prior condition that was significantly different from chance (Figure 2c, 0.591; t(23)=18.47, p<0.001), but significantly lower than for the positive prior condition (t(23)=2.316, p<0.05). The Partial Explicit and Explicit groups performed similarly, with no significant difference between the ability to apply the priors associated with the two colors (p = 0.889), even though participants in the Partial Explicit group had minimal awareness of how stimulus color was associated with orientation probability.

**Figure 2:**
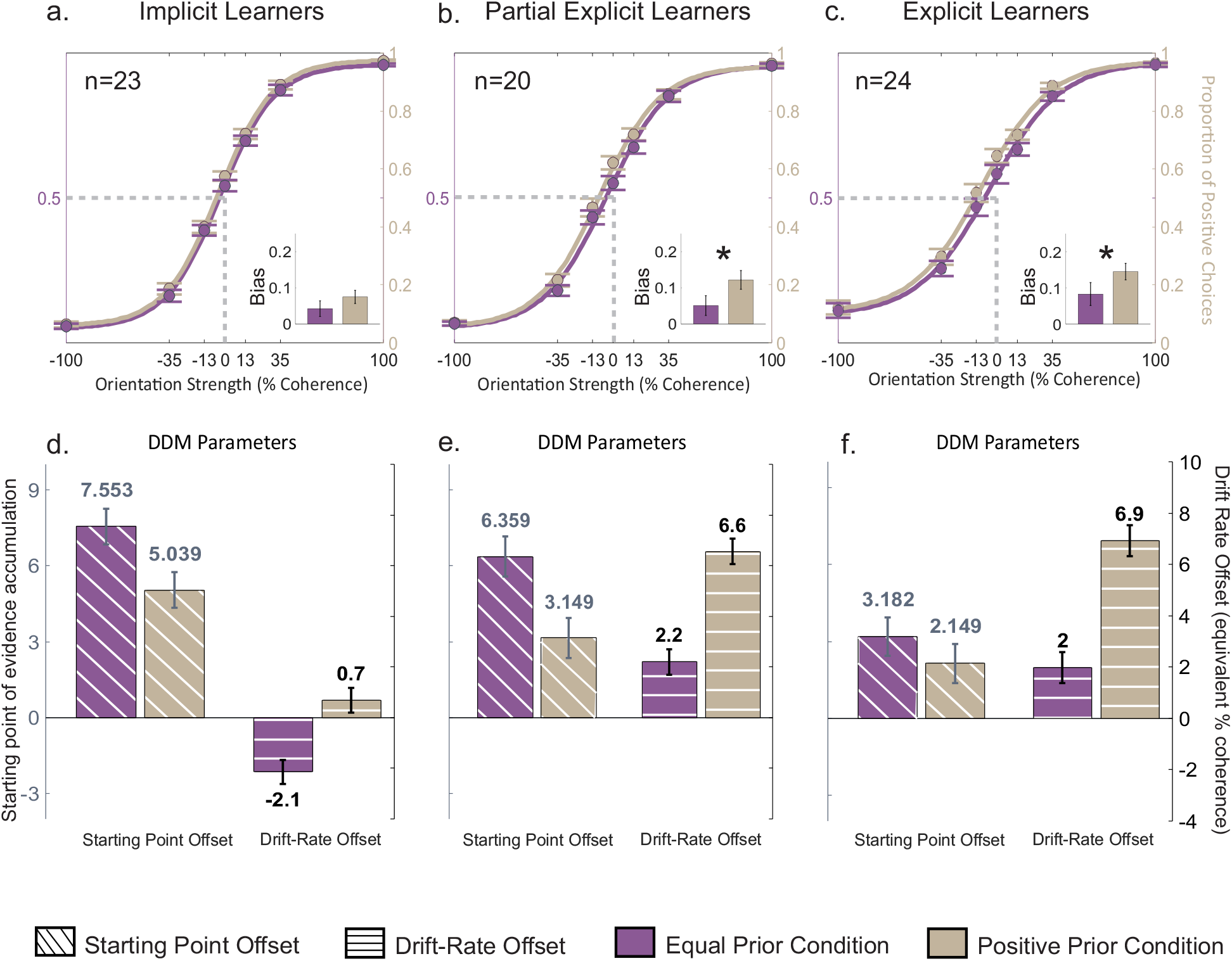
Participants learn multiple priors implicitly and explicitly. a. Proportion of choices plotted against Glass pattern coherence for 23 participants. Filled lilac circles and error bars show the mean and SE for each coherence across all participants who reported no difference in orientation frequency (Implicit Learners) for the equal prior condition (50% positive and 50% negative). The lilac solid lines show the best fit logistic function (see methods). The khaki filled circles and lines show the same for the positive prior condition in which the more frequent orientation was positive, referred to as positive prior condition (75% positive and 25% negative). The inset shows the bias parameter for equal (lilac) and positive (khaki) priors represented as deviation of the proportion of positive choices at 0% coherence from unbiased condition (0.5). b. Same as in a for the participants who reported a difference in orientation frequency regardless of color or for only one color (Partially Explicit Learners; n = 20). c. Same as in a for the participants who were informed of the prior conditions (Explicit Learners; n = 24).d. Parameter estimates from the DDM fits for the equal prior condition (lilac) and the positive prior condition (khaki) for all Implicit Learners. Diagonal hatched bars show the mean and 95% confidence intervals (CI) of the starting point offset and horizontal hatched bars show mean and 95% confidence interval for the drift-rate offset (equivalent % coherence). e. Same as in d for the Partially Explicit Learners. f. Same as in d for the Explicit Learners.

### Implicit and Explicit Learners Use Different Mechanisms to Apply Decision Biases

Using the DDM, we previously observed that healthy participants adjust both the starting point of evidence accumulation and the drift rate offset to express decision-making biases, and that it was the change in the drift rate offset that provided feature specificity to the bias^26^. However, in that study we did not know whether the healthy participants were aware of the feature-prior association or whether awareness would be associated with the use of different mechanisms to implement the bias. Here, we assessed whether the mechanism underlying the expression of the bias differed for the different types of Learners, Implicit, Partially Explicit and Explicit.

We fitted two feature-specific DDMs to the second-half of the experimental session data independently (600 trials), allowing only the starting point and the drift rate offset to vary between the two features (Table 1). Figure 2d-f shows the mean parameter estimates and confidence intervals of the parameter estimates for the starting point and drift rate offsets for both prior conditions. For the Implicit Learners, the starting point shifted toward the more frequent orientation for both the equal and the positive prior trial s. The shift in starting point was similar for the two conditions (Equal Prior-estimate: 7.6%, S.E.: 0.7%, Positive Priors-estimate: 5%, S.E.: 0.7%). The change in starting point was numerically larger for the equal prior condition compared to the positive prior condition, but this was counteracted by an opposite change in the drift rate offset for equal prior trials (estimate: -2.1%, S.E.: 0.5%). The drift rate offset for the positive prior condition was in the same direction as the more frequent orientation but with a small magnitude (estimate: 0.7%, S.E.: 0.5%). In summary, for the Implicit Learners, there was a significant increase in starting point for both prior conditions accompanied by a slight decrease in drift offset when priors were equal, leading to a bias in decision-making that was undifferentiated by stimulus color.

For the Partially Explicit Learners, we observed similar changes in the starting point of evidence accumulation and quantitatively larger changes in the drift rate offset compared to the Implicit Learners (Figure 2e). The starting point of evidence accumulation for equal prior trials was 6.4% (S.E.: 0.8%) and 3.1% for the positive prior trials (S.E.: 0.8%). Similarly, the drift rate offset was positive for both prior conditions but in contrast with the starting point, the drift rate was much higher for the positive prior condition (6.6%, S.E.: 0.5%) compared to the equal prior condition (2.2%, S.E.: 0.5%). Thus, both Implicit and Partially Explicit Learners expressed a decision bias by changing the starting point, however, only Partial Explicit Learners developed a sensitivity to priors associated with stimulus colors that appears to be mediated primarily by changes in the drift rate offset.

We found that the Explicit Learners also showed an overall bias but only a numerically greater feature specific sensitivity to the priors unlike the Partially Explicit Learners (Figure 2b and c). The starting point offsets in the Explicit Learners were similar for both priors (Equal Prior: 3.2%, S.E.: 0.8%; Positive Prior: 2.1%, S.E.: 0.8; Figure 2f). Similar to Partial Explicit Learners, the Explicit Learners also adjusted their drift rate offset asymmetrically to express the bias (Equal Prior: 2.0%, S.E.: 0.6%; Positive Prior: 6.9%, S.E.: 0.6%; Figure 2f). Hence, similar to the Partial Explicit group, the feature-specific biases were primarily driven by drift rate offset in the Explicit group.

To see the effect of starting point offset or drift rate offset individually, we simulated the DDM model using optimized parameters with minor modifications. To identify the contribution of the starting point offset, we manually kept the drift offset at zero and measured biases at 0% coherence for both prior conditions. Similarly, to identify the contribution of the drift rate offset to the bias, we simulated the model with optimized parameters and manually set the start point offset to zero. We observed that with the full model the biases in the Implicit group, Partial Explicit group and Explicit group were: [Equal Prior, Positive Prior]: [0.531, 0.555]; [0.567, 0.602]; [0.585, 0.622] respectively. When we looked at only the starting point’s contribution, we found simulated biases as follows: [Equal Prior, Positive Prior]: [0.555, 0.547]; [0.543, 0.531]; [0.567, 0.558] for the Implicit group, Partial Explicit group and Explicit group respectively. When only the drift offset was kept in the model, we found the biases for the Implicit, Partially and Explicit and Explicit groups to be: [Equal Prior, Positive Prior]: [0.484, 0.524]; [0.534, 0.586]; [0.567, 0.610]. As we can see from the above analysis, the start point contributes toward an increase in the overall bias of the choice behavior and the drift offsets contribute toward the color specificity in all three groups including the Implicit group. However, the drift rate offsets in the Implicit group were smaller than start point offset leading to a lack of a significant color-specific bias.

In Experiment 1, participants were able to learn a Bayesian prior implicitly based on different base rates of orientation and could apply this prior in a perceptual decision-making task. Participants were also able to apply priors differentially depending on stimulus color even if they were unaware that the colors were associated with different base rates. Implicit knowledge of the general prior, that one of the orientations occurred more frequently, was captured by a start point offset in the DDM. Explicit knowledge of the color-specific priors was captured mainly by changes in drift rate offset in the DDM. Participants who gained some knowledge of the frequency of the orientations across the task but did not become aware of the color-specific priors also showed differences in drift rate offset for the two conditions.

### Experiment 2-Learning Feature -Specific Priors in the Absence of a General Prior

The design of Experiment 1, with an equal base rate prior for one stimulus color (50% for each orientation) and a positive prior for the other (75% for one orientation) led to a general prior of 67% towards the positive orientation across all stimuli. Participants in all groups generalized across the stimulus colors to some extent with biased responses toward the positive prior. Thus, it was difficult to tease apart learning feature-specific and general prior knowledge in the results. To examine learning of feature-specific priors in the absence of a task general prior, we conducted a second experiment in which the base rate orientation of the two stimulus colors were in opposite directions. So, for example, the Glass patterns of one color occurred with 75% of trials with a positive orientation and 25% of the trials with a negative orientation, whereas the Glass patterns of the other color occurred with a positive orientation on 25% of the trials and a negative orientation on 75% of the trials (Fig 1c). This arrangement ensured that the overall positive and negative orientation probabilities remained constant.

We analyzed the data for a different group of 41 participants in Experiment 2, all with informed consent and using procedures approved by the UCLA IRB. 21 participants were assigned to the Implicit Learner group as they were not told about the differences in base rates. None of these participants reported noticing base rate differences on the questionnaire given after the session. 20 participants were assigned to the Explicit group, and were informed of the different base rates for the two colors. Figure 3a shows that even though Implicit Learners were unaware of the priors, they showed a feature-specific bias. The mean bias for the positive prior trials was 0.540 and for the negative prior trials was 0.423 and this difference in biases were statistically significant (Figure 3a: t 2.68, p<0.001). Using the DDM, we found that both the starting point and the drift rate offsets were asymmetric with the task feature i.e., both were negative for negative prior and non-negative for the positive prior condition. The change in starting point for the negative priors was -1.6% (S.E.: 0.8%; Figure 2c) and for positive priors was 0% (S.E.: 0.8%; Figure 2c). The drift rate offset parameter, like the starting point, changed such that the negative prior had a negative drift rate offset and the positive prior had positive drift rate offset: Negative Prior: 3.7%, S.E.: 0.6%; Positive Prior: 4.7, S.E.: 0.6%; Figure 3c). These results are consistent with Experiment 1 in that an implicitly learned feature-selective prior biases decision-making through both starting point and drift rate offset with a dominant effect from the drift rate offset.

**Figure 3:**
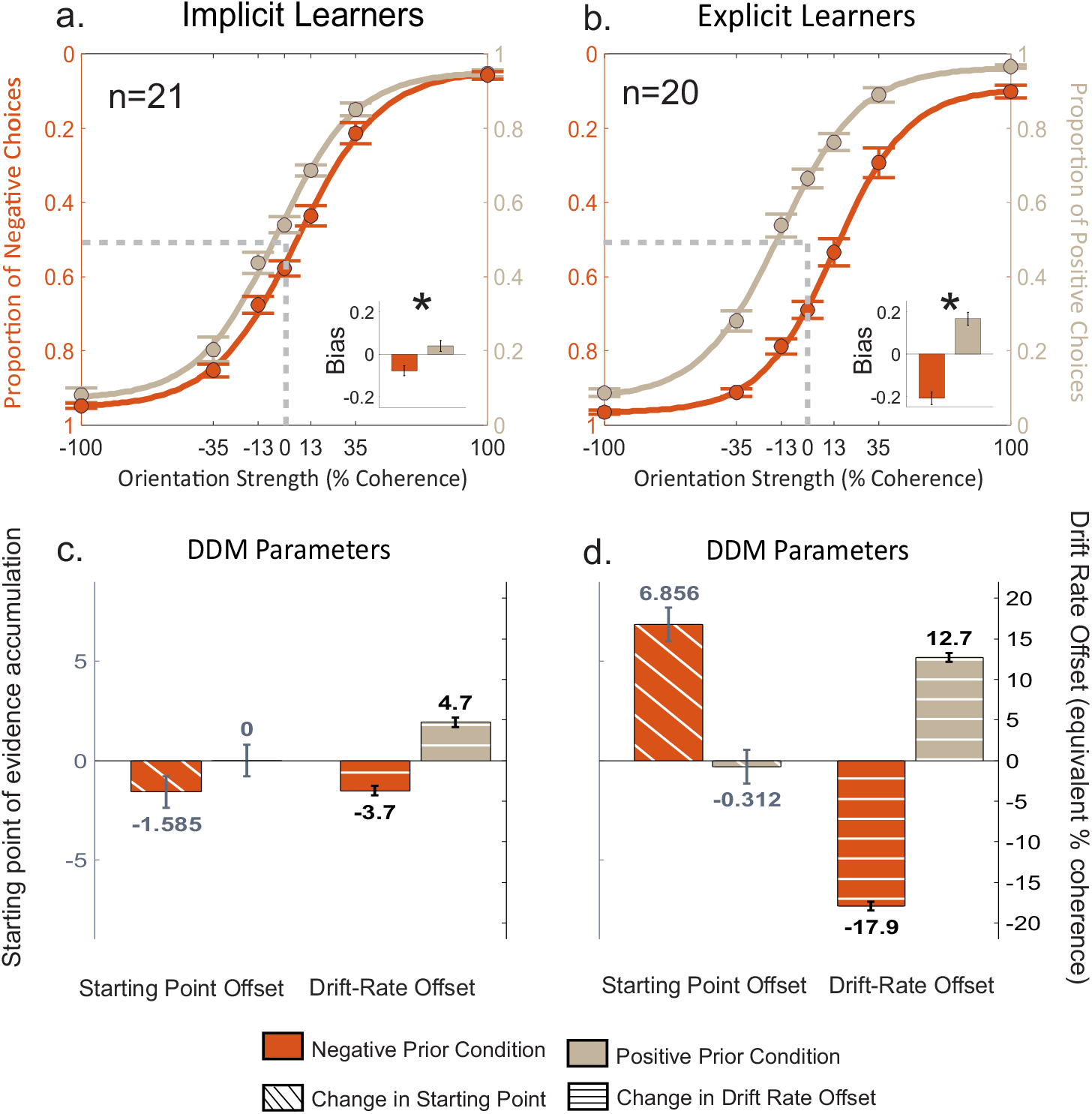
Participants learn and use stimulus specific orientation priors that bias perceptual decisions. a. Proportion of choices plotted against Glass pattern coherence from 21 participants. Filled orange circles and error bars show the mean and one SE for each coherence across all participants who reported no difference in orientation frequency (Implicit Learners) unequal prior condition in which the more frequent orientation was negative referred to as Negative Prior condition (75% negative and 25% positive). The orange lines show the best fit logistic function (see methods). The khaki filled circles and lines show the same for the unequal prior condition in which the more frequent orientation was positive referred to as Positive Prior condition (25% negative and 75% positive). The inset shows the bias parameter for negative (orange) and positive (khaki) priors. It represents the deviation of proportion of positive choices for 0% coherence from the unbiased condition (0.5). b. Same as in a for the participants who reported a difference in orientation frequency (Explicit Learners; n = 20). c. Parameter estimates from the DDM fits for the negative (orange) and positive (khaki) prior conditions for all implicit learners. Diagonal hatched bars show the mean and 95% CI of the parameter estimates for the starting point offset and horizontal hatched bars shows the mean and 95% CI of the parameter estimate for the drift-rate offset (in equivalent % coherence). d. Same as in c for the Explicit Learners.

Participants in the Explicit Group showed robust expression of feature-specific biases. The mean bias measured for the 0% coherence, positive prior trials was 0.662 and for the negative trials was 0.292 and these differences were significantly different (Figure 3b; t(19)=7.13, p<0.001) and the difference in bias for the two colors was significantly different from those observed in the Implicit Learners (t(39)=3.782, p<0.001). The modeling results showed that the Explicit Group used drift rate offsets to express stimulus feature-prior associations, and these changes were much larger than those obtained in the Implicit Learners. In the Explicit group, the starting point offset was actually in the opposite direction to the priors. The starting point estimates for the Explicit group were, Negative Prior: 6.9%, S.E.: 0.8%; Positive Prior: 0.3%, S.E.: 0.8%; (Figure 3d). Similar to the Implicit Learners, the Explicit group also showed feature-specific changes in the drift rate offset and these changes were much more prominent: Negative Prior: -17.9%, S.E.: 0.6%; Positive Prior: 12.7%, S.E.: 0.5%; (Figure 3f). This result suggests that the change in starting point for the negative prior condition could be a compensatory mechanism for a high change in the drift rate offset as reported previously^26^. These results indicate that according to the DDM, people implement explicit priors through the enhancement of drift rate toward the decision boundaries.

In Experiment 2, we wanted to measure the individual contribution of starting point and drift rate offset when there were priors that were stimulus specific with no overall prior. We observed that with the full model the biases in Implicit Learners and the Explicit group were: [Negative Prior, Positive Prior]: [0.448, 0.550]; [0.318, 0.646] respectively. When we looked at only the starting point’s contribution, we found simulated biases as follows: [Negative Prior, Positive Prior]: [0.482, 0.507]; [0.500 0.516] for the Implicit and Explicit groups respectively. When the drift offset was kept in the model, we found the biases for the Implicit and Explicit groups to be: [Negative Prior, Positive Prior]: [0.457, 0.550]; [0.287, 0.648]. Similar to Experiment 1, the drift rate offset contributes to the stimulus specific biases. The starting point had a smaller contribution that was actually compensatory for the large drift rate offsets in the Explicit group.

Based on the results of the two experiments, implicit learning of base rate information can occur in both a general and a feature-specific manner, and this knowledge appears to influence decision-making through the more efficient accumulation of sensory evidence consistent with the priors. When participants are aware of the base rate differences, there are greater increases in the drift rate offset. Starting point offsets can also contribute to the stimulus specificity either by increasing the bias of smaller changes due to drift rate offset, as in the Implicit Learners or by counteracting the large changes in drift rate offset seen in the Explicit group. Thus, starting point may be adjusted to both bias decision-making implicitly and possibly to compensate for large drift rate increases that implement explicit knowledge of priors.

## DISCUSSION

We assessed the ability of participants to learn base -rate priors implicitly and apply them in a perceptual decision-making task. In both experiments, participants acquired a bias in an orientation judgment task that was apparent when participants judged items with zero coherence. Participants who did not report awareness of the priors nevertheless exhibited evidence of bias in their decisions about the orientations of the stimulus. In Experiment 1, two different kinds of priors were present; an overall prior based on the frequency of the different orientations across all stimuli and feature-specific priors that were specific to stimulus color. We found that both types of priors could be learned implicitly. In the group that reported no explicit knowledge of the different priors of the two orientations (Implicit Learners), there was still a significant bias toward the more frequent orientation across all stimuli, which was also seen in participants who were informed of this prior or became aware of it during testing. Implicit learning of feature-specific priors was also seen in those participants who became aware of only the general difference in frequency between the orientations (Partial Explicit Learners). In this group, explicit knowledge of the structure of the task lagged behind implicit knowledge, in that they did not become aware of differences in priors for the two colors despite applying the differential priors in their choice behavior. In Experiment 2, the two orientations were equally likely overall and the priors for the two colors were more distinct than in Experiment 1. Here, we also found implicit learning of feature-specific priors.

Our results are consistent with the recent study by Rungratsameetaweemana et al., 2019^32^ in which the ability of patients with amnesia following hippocampal damage to acquire a prior in a motion direction judgment task was measured. The sensitivity to prior information was similar in amnesic patients and control participants, supporting the idea that explicit learning of the prior does not benefit performance. We assessed awareness of priors in participants after the session to determine the level of explicit knowledge gained during the task. We also demonstrated that different priors specific to different sets of stimuli based on color could be learned implicitly. These data suggest that changes in perceptual processing can occur based on implicit knowledge of priors rather than only in overall response bias.

Another goal of the present study was to compare the contribution of explicit and implicit knowledge of priors to decision-making. Using the DDM, we found that explicit and implicit knowledge of the priors appeared to influence decisions differently. Substantial drift rate modulation was associated with knowledge of the color-specific priors and was much more robust in groups with explicit knowledge than in groups with only implicit knowledge. The influence of starting point on decision-making was confined to implicitly learned knowledge of the priors. The Implicit group in Experiment 1, who learned to apply a bias towards the more frequent orientation, showed a significant starting point offset in the positive direction for both colors. In Experiment 2, where the priors were in equal and opposite directions, the bias in decision-making occurred as a result of significant changes primarily in drift rate offset for stimuli of the different colors. The difference in the drift rate offsets for the two stimulus colors was relatively modest for the Implicit group. In contrast, in the group that was aware of the different priors, there were substantial increases in drift rate offset towards the positive and negative direction for the two colors of stimuli. In Implicit Learners, the starting point supported further increases in bias by offsetting in the direction of the prior for one of the colors. In contrast, in the Explicit group the high change in drift offset rate was counteracted in part by a start point offset in the opposite direction for one of the priors. These results suggest that explicit knowledge of the priors leads to faster accumulation of evidence in the direction of the prior across coherences. Implicit knowledge of stimulus specific priors appears to also affect drift rate offset. Start point offsets appeared to reflect implicit knowledge of priors, and may even compensate for large drift rate offset changes implemented by explicit knowledge of priors.

The present work shows that people can implicitly learn base-rate differences and apply this knowledge in a perceptual-decision making task. While classic work in decision-making shows that participants often ignore explicit base-rates^33^, base-rates may have a greater impact when they are learned through experience. An implicit learning mechanism may be engaged with experiential learning that may lead to knowledge that is more effective in influencing judgments than explicit knowledge ^34^. We argue that implicit learning of general base-rate information shifts the starting point of evidence accumulation toward the more frequent outcome. Consistent with this idea, local fluctuations in base rates influence decision-making through starting point modulation, even when these local fluctuations do not reflect the explicit priors for the entire task^35^. Thus, starting point, perhaps reflecting biasing activity in neural representations of alternatives, may be modulated fairly automatically by reinforcement history.

Additionally, we also observed changes in the drift rate offset associated with implementing feature-specific biases in decision-making. When participants had knowledge that there were different priors that depended on stimulus color, there was a significant modulation of drift rate, indicating that awareness of feature-specific priors led to greater efficiency in accumulating perceptual evidence in support of the priors. Drift rate changes were also observed despite lack of awareness that the different stimulus colors were associated with different priors in both experiments. It thus appears that implicit knowledge of stimulus specific priors enhances the perceptual processes contributing to decision-making.

Modulation of starting point and diffusion rate are likely supported by different brain regions in the service of decision-making. Starting point offsets may reflect increased activity in neural representations corresponding to more frequent alternatives. In the seminal work of Basso & Wurt, 1998^36^ and Platt & Glimcher, 1999^37^, monkeys performing a two-alternative forced-choice task showed increased pre-trial activity in SC and area LIP representation for the alternative that was cued to be more likely. In Experiment 1, a similar increase in activity in representations corresponding to the more frequent alternative may have occurred. This increase may have occurred very early in the trial, before the integration of evidence about orientation. In the present study, subjects were not cued with base-rate priors but were able to acquire them implicitly through experience. Thus, it appears that modulation of start point does not require awareness of priors. Perugini et al., 2016^26^ found that patients with Parkinson’s disease are impaired at adjusting starting point based on priors, suggesting that basal ganglia might also be involved in changes in starting point and adjusting decision threshold generally ^38^. Future work may clarify whether modulation of starting point in decision-making relies on similar neural mechanisms for explicitly cued and implicitly learned priors.

The adjustment of drift rate offset in the present study may reflect enhanced processing of elements of the display consistent with the prior. Adjustment of drift rate has been shown to occur in decision-making paradigms in which there is variability of signal to noise across trials^39^; as variability of difficulty across trials increased, fitting the DDM to data relies more heavily on adjustments to the drift rate^40^. In the present study, where trials varied substantially in terms of difficulty, it is therefore not surprising that drift rates were offset resulting in decision biases. Although present in the Implicit group in Experiment 2, this finding was much more striking in subjects who gained partial knowledge of the base rates or were informed of the different priors at the onset of the task. One possibility is that drift rate offsets reflect top-down attentional effects that enhance gain in perceptual or decision-making regions. This may result from phasic activity in the locus coeruleus that is driven by orbitofrontal and anterior cingulate inputs. This increase in gain may be applied to pools of neurons processing one orientation or the other, or representing one alternative or the other, depending on the color of the stimulus.

In sum, our results demonstrate that knowledge about priors can be acquired implicitly and that this knowledge can be specific to stimuli with different features within a stimulus set. Within the DDM framework, the implicit acquisition of overall bias seems to be implemented through a change in starting point offset. However, feature-specific bias is primarily implemented through changing the rate of evidence accumulation. Future studies can help to identify the brain regions involved in implementation of general and feature-specific biases, and whether implicitly acquired biases are supported by different neural mechanisms than explicit biases.

## METHODS

### Participants

We recruited 140 people for this study. Experiment 1 included 89 participants (23 male and 66 female) and Experiment 2 included 51 participants (13 male and 38 female). All participants were right-handed UCLA students with age ranging from 18 to 30 years. None of the participants had active medical, neurological, or psychiatric diagnosis, and were not taking any chronic medication that could affect sensory processing, movement, or cognition. All participants had normal/corrected to normal vision. All the participants were tested for colorblindness using an online version of the Ishihara color blindness test. All the participants gave written informed consent as per the guidelines of the Institutional Review Board of the University of California and received course credit for their participation. In each experiment, subjects were assigned to either an Explicit group, in which they were informed of the different orientation probabilities for red and green stimuli or an Implicit group, in which they were not informed. All subjects were given instructions by the experimenter and completed at least 50 practice trials of stimuli that were 100% coherent with 85-90% accuracy to familiarize themselves with the task before beginning.

#### Visual Stimuli

Dynamic translation Glass patterns were used for this study, which was developed previously^26^. This task uses two identical dot patterns, one of which is translated in position with respect to other. When these patterns are superimposed, multiple correlated dot pairs are formed, which results in a strong perception of the orientation of dots ^24^. The strength of this orientation can be varied using the number of dot-pairs formed. For both experiments, we used 0%, 13%, 35% and 100% coherence, referring to the percentage of correlated dot pairs. In the 100% coherence condition, all dots pairs are correlated, and the orientation of the stimulus (right or left) is readily apparent. In contrast, the 0% coherence condition does not have any dot pairs which are correlated and thus subjects have no perceptual information upon which to base their orientation decision. The stimuli were made dynamic by generating 30 frames of such images and showing them with a frequency of 85 frames/ sec.

#### Experimental Paradigm

Each trial began with the appearance of a centrally-located white fixation box. Subjects were asked to keep looking at the fixation box. After fixation ranging 1000ms ± 200ms, two target boxes appeared on the screen at left-upper and right-upper positions. Immediately after that, a Glass pattern with one of the coherences mentioned above was shown to the subjects. In each trial, the glass pattern was shown either in red (luminance 0.6 cd/m^2^) or green (luminance 0.54 cd/m^2^), which was randomized with equal probability. Subjects had to gather the information about the orientation of the stimulus and record their response as quickly as possible by pressing a key on the keyboard. If they judged the stimulus to be oriented to the left, they pressed the ‘O’ key, and if they judged it to be oriented to the right, they pressed the ‘P’ key. The Subjects were given 3 seconds after the onset of the Glass pattern to decide the orientation. If they did not respond within 3 seconds, the trial was aborted. If they chose the correct orientation, the feedback was given as a high tone by the computer. For both experiments, each subject performed at least 1200 trials.

For Experiment 1, one color of Glass pattern was unbiased regarding orientation (both left and right were equally likely) and for the other color, the orientation occurred 75% in one orientation and 25% in the other. For Experiment 2, both the red and green patterns were biased, with one orientation occurring 75% of the time and the other 25% of the time, and the opposite probabilities occurred for the other color. Thus, the two orientations (left and right) occurred equally often across the stimulus set. The assignment of colors to conditions was counterbalanced across subjects for both experiments and later converted to positive orientation and negative orientation. In this way, we were able to examine whether both general biases (one outcome is more likely to occur in the task) and feature-specific biases (red and green stimuli have different base rates) can be learned in the absence of awareness.

Following the completion of the task, participants’ awareness of the priors was assessed using a follow-up questionnaire. The questionnaire asked participants if they thought that each colored stimulus (red and green) was equally distributed between two orientations. Based on their response, participants were divided into two categories: 1) Implicit Learners and 2) Partial Explicit Learners. Participants who responded that both orientations occurred equally for each color, i.e., there was no bias, were categorized as Implicit Learners. Participants who identified the prior orientation of only one stimulus category correctly were categorized as Partial Explicit Learners. In the Explicit group, participants were informed about the feature-prior associations at the start of the experiment.

#### Data Analysis

##### Behavior

The reaction time (RT) was defined as the duration between onset of Glass pattern stimulus and response time (keypress) onset. All trials with RT less than 200 ms were excluded from the final analysis due to the possibility of anticipatory trials. Additionally, all participants whose performance at 100% coherence was below 90% accuracy were removed from the final analysis (n=32, across all five groups). Hence, data was analyzed for 67 participants in Experiment 1 and 41 participants in Experiment 2.

Next, a psychometric function of accuracy with respect to coherency was calculated. This psychometric function was fitted using the following logistic function of the form:

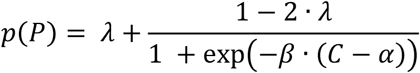

Where *p*(*P*) is the proportion of choices towards more frequent orientation, i.e., for 75 -50 condition with positive prior, *p*(*P*) is the proportion of positive choices; *C* is dot pair coherence; *α* and *β* are free to model parameters and are determined by the maximum likelihood method. They also represent response bias and sensitivity to stimulus, respectively. *λ* represents the lapse rate, which is the difference between asymptotic function and perfect performance. It indicates the transient lapse in attention during the experiment ^41^. As 0% coherence is an utterly ambiguous stimulus, subjects’ biases can be assessed at that coherence condition. The use of priors for each stimulus color was calculated as the proportion of choices towards the more frequent orientation at 0% coherence level per subject. The statistical analysis was carried out using pairwise t-test.

##### Modeling

Drift Diffusion Model: One well-known example of the class of evidence accumulation models is the Drift Diffusion Model (DDM). This model explains both, the reaction time and the choice of every trial. We used one of the variants of this model with a collapsing boundary, i.e., as time passes, participants require less accumulation of evidence to make a decision to account for urgency. We fit the DDM model with eleven parameters on pooled data across all participants in a given condition of the experiment. These eleven parameters were: the non-decision time, starting point of accumulation, proportionality factor between coherence and drift rate, the diffusion coefficient, a scaling parameter for collapsing bound, delay in collapsing bound, drift rate offset, proportion of uninformed positive choices, proportion of uninformed negative choices, mean and standard deviation of reaction time for uninformed choices. The biases can be implemented in the DDM-like model by either changing the starting point of evidence accumulation (Figure 1e) or by changing the rate of evidence accumulation across all coherences i.e., drift rate offset. This drift rate offset can also be interpreted as the apparent coherence during 0% coherence stimulus. (Figure 1f). Each of these methods predict slightly different behavior. The change in starting point predicts an asymmetric change in RT between positive and negative choices. On the other hand, a drift rate offset predicts a similar change in RT for positive and negative choices, such that zero coherence trials are no longer the slowest ones. To assess these two mechanisms, we fitted two DDM models, one for each stimulus color, such that five of the model parameters, namely, non-decision time, proportionality factor, the diffusion coefficient, scaling and delay of collapsing bound were shared by both models. The other two parameters, starting point and drift rate offset, were allowed to differ between both models, thus allowing us to assess which parameter change or combination of changes could better explain the behavioral data. The DDM model was fitted to only the second-half of the session for each subject, the rationale being that during the first half of the session subjects are still learning these priors and decision behavior will be more stable during the second half of the session.

Under drift-diffusion models, a decision-making process is initiated at the starting point and continues to accumulate noisy evidence, thus continues to drift until it reaches one of the decision bounds. The decision bounds are defined as:

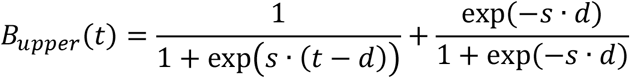

Where, *s* is a decay parameter of the boundary, *d* is a delay parameter to start the decay and *t* is time. *B*_*lower*_ is defined as *B*_*upper*_. Hence at *t* =0, the bounds are at ±1 and decay towards zero as time progresses. The drift rate (rate of accumulation) *r* is defined as

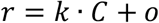

where C is the signed coherence of the stimulus such that positive C means coherence towards positive orientation, and o is the offset in the drift rate. The reaction time can be obtained by

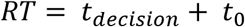

where *t*_*decision*_ is the time required to cross a decision bound, and t_0_ is the non-decision time to account for the sensory and motor processing delays.

The traditional DDM model does not account for lapses in the data. To account for this, we added four parameters to our model such that on random proportions of trials subjects provide an uninformed response with a RT derived from an unknown Gaussian distribution. The mean and the standard deviation of the Gaussian distribution are estimated as model parameters from the data. Hence, the additional four parameters are: proportion of uninformed positive choices, proportion of uninformed negative choices, mean and standard deviation of reaction time for uninformed choices. The proportion of uninformed positive and negative choices were fitted separately for each color.

All the behavioral data analyses and modeling was performed using MATLAB. The DDM model was implemented from the Stochastic Integration Modeling Toolbox for MATLAB which was developed by J. Ditterich (https://www.peractionlab.org/software).

The parameter optimization was carried out using maximum likelihood estimation accounting for both reaction time and choice on each trial. We divided the data into 16 subsets based on the coherence (4x), direction (2x), and choice (2x). The likelihood of choice and RT (using QMLE method on normalized histogram of RT distribution) for each subset was calculated independently and log summed at the end for all subsets.

The reaction time likelihood was calculated using modified Quantified Maximum Likelihood Estimate (QMLE)^42^. To get better resolution of estimated RT distribution, we interpolated RT distribution from model and data to 1ms resolution. Next, we computed the log likelihood from model for each millisecond and took its weighted sum based on RT distribution from data.

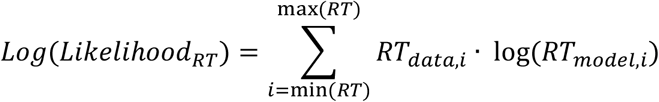

Once RT likelihood was calculated for all 16 subset of conditions, we calculated a weighted sum based on the number of trials in each subset. The choice likelihood was calculated using the binomial probability equation ^43^, as this was a 2AFC task:

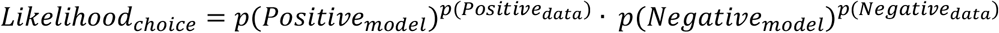

Similar to RT likelihood, choice likelihoods were log summed across all 16 subsets of conditions, and, finally, both RT log likelihood and choice log likelihood were summed. This total likelihood was later optimized using MATLAB functions.

We used a combination of a multidimensional simplex approach (“fminseach” in MATLAB Optimization Toolbox) and a pattern search algorithm (“patternsearch” in MATLAB Global Optimization Toolbox) to find the global optima. Standard errors of the estimated parameters were obtained using 1-dimensional local Gaussian approximation of the likelihood function (*L*) around the optimal value *p*_*opt*_ of each parameter p:

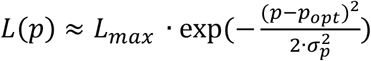

