## Supplemental Data for "Implicit and explicit learning of Bayesian priors differently impacts bias during perceptual decision-making"

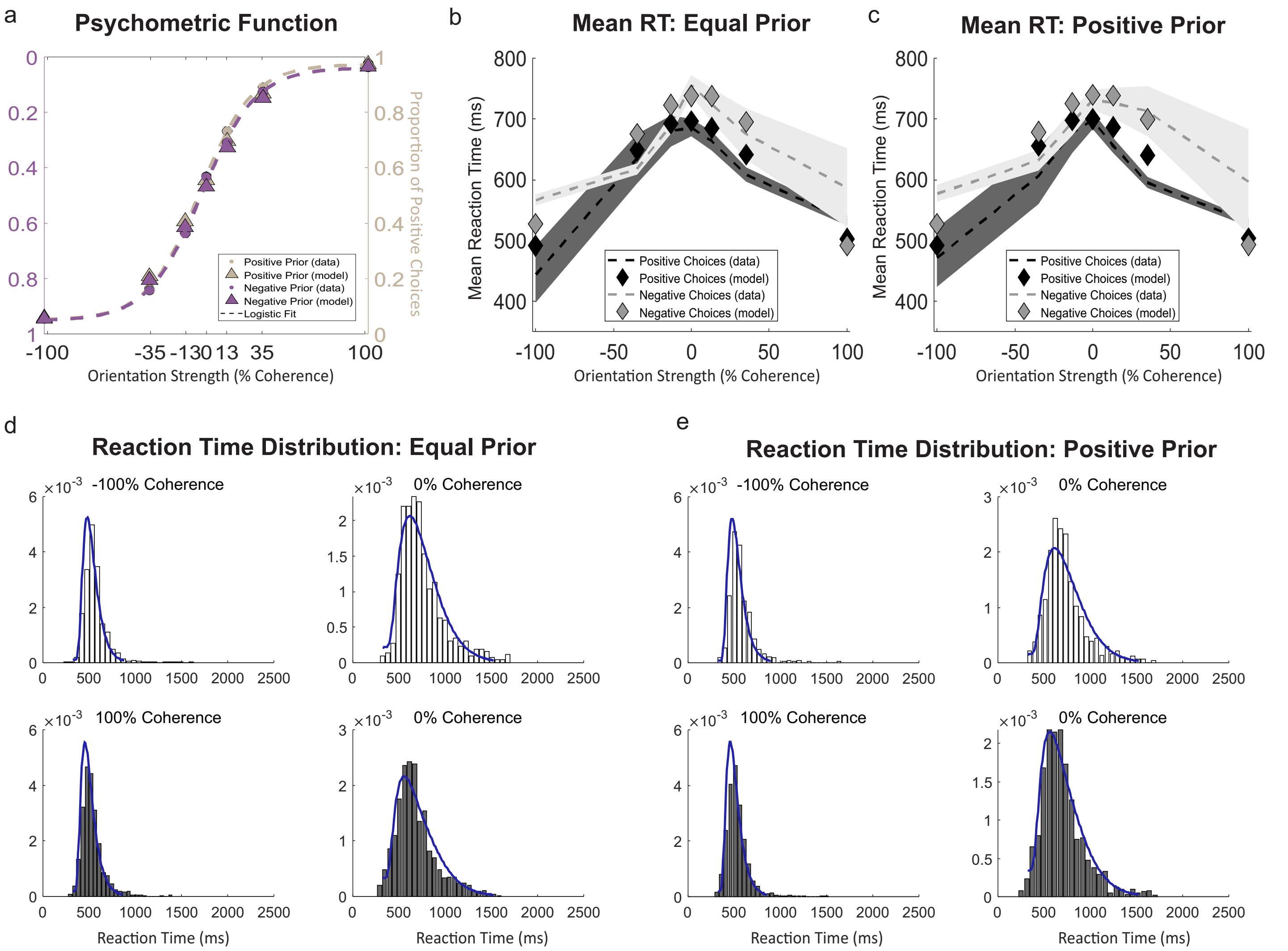


Figure S1. Model fits for choices and reaction times for Implicit condition in Experiment 1.

a. Psychometric function showing proportion of positive choices as a function of coherence for Equal prior condition (*lilac*) and Positive prior condition (*khaki*). Circles shows the data and the triangles show the model predictions and the dashed line shows the maximum likelihood logistic fit to the data.

b. Mean reaction time plot as a function of stimulus coherence for Equal prior condition. The gray color shows the mean RT for negative choices and dark color shows mean RT for positive choices. Dashed line is the mean reaction time and the shaded area shows the 95% confidence interval. The model prediction of mean RT is shown by the diamonds.

c. Same as b. for Positive Prior condition.

d. Reaction Time distributions under Equal prior condition for i) 100% coherence in negative direction and negative choices (gray, upper left), ii) 0% coherence and negative choices (gray, upper right), iii) 100% coherence in positive direction and positive choices (dark, lower left), and iv) 0% coherence and positive choices (dark, lower right). The histogram shows the data and blue traces show the model prediction.

e. Same as d. for Positive prior condition.

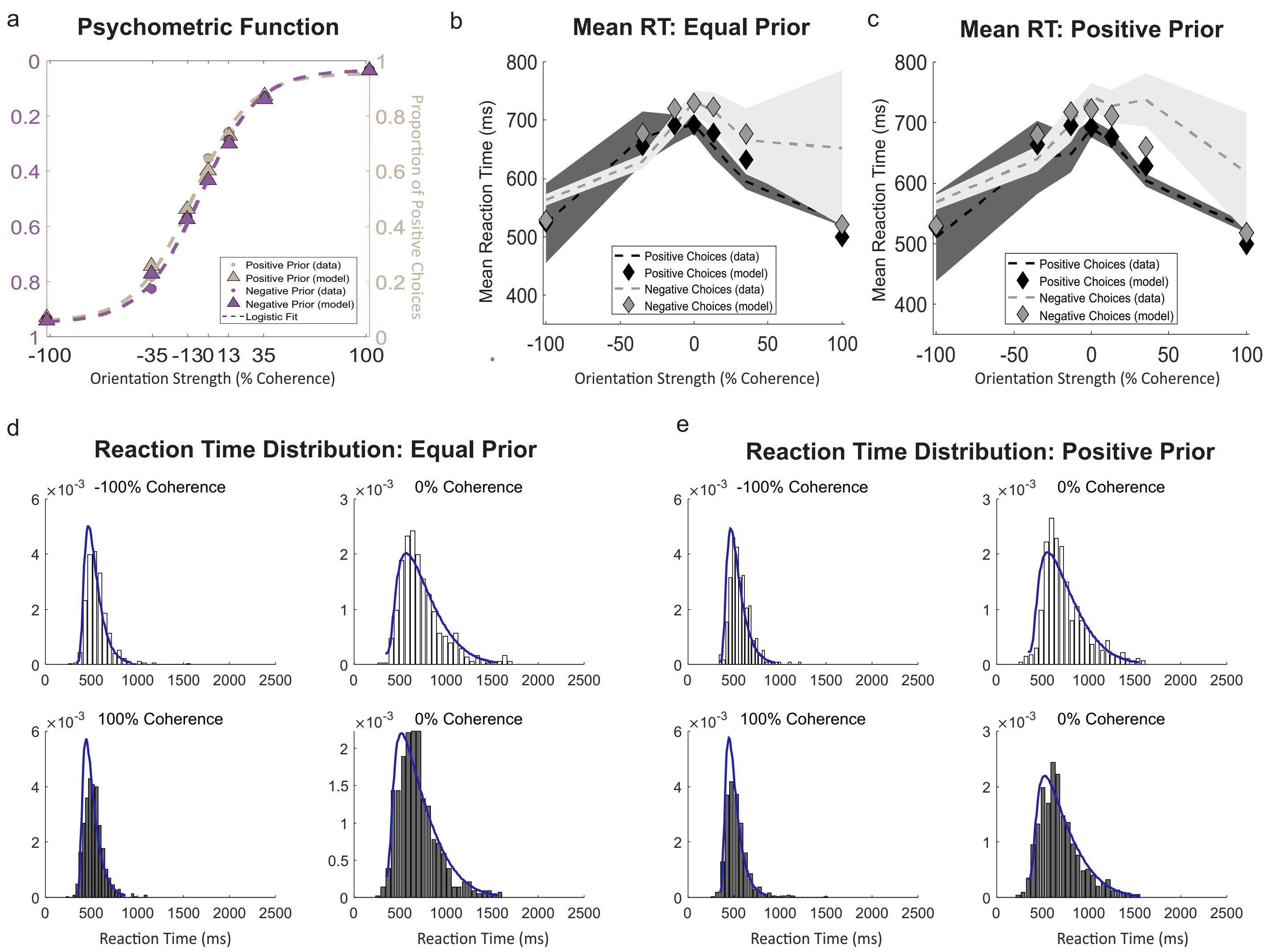


Figure S2. Model fits for choices and reaction times for Partial Explicit condition in Experiment 1.
a. Psychometric function showing proportion of positive choices as a function of coherence for Equal prior condition (*lilac*) and Positive prior condition (*khaki*). Circles shows the data and the triangles show the model predictions and the dashed line shows the maximum likelihood logistic fit to the data.
b. Mean reaction time plot as a function of stimulus coherence for Equal prior condition. The gray color shows the mean RT for negative choices and dark color shows mean RT for positive choices. Dashed line is the mean reaction time and the shaded area shows the 95% confidence interval. The model prediction of mean RT is shown by the diamonds.
c. Same as b. for Positive Prior condition.
d. Reaction Time distributions under Equal prior condition for i) 100% coherence in negative direction and negative choices (gray, upper left), ii) 0% coherence and negative choices (gray, upper right), iii) 100% coherence in positive direction and positive choices (dark, lower left), and iv) 0% coherence and positive choices (dark, lower right). The histogram shows the data and blue traces show the model prediction.
e. Same as d. for Positive prior condition.

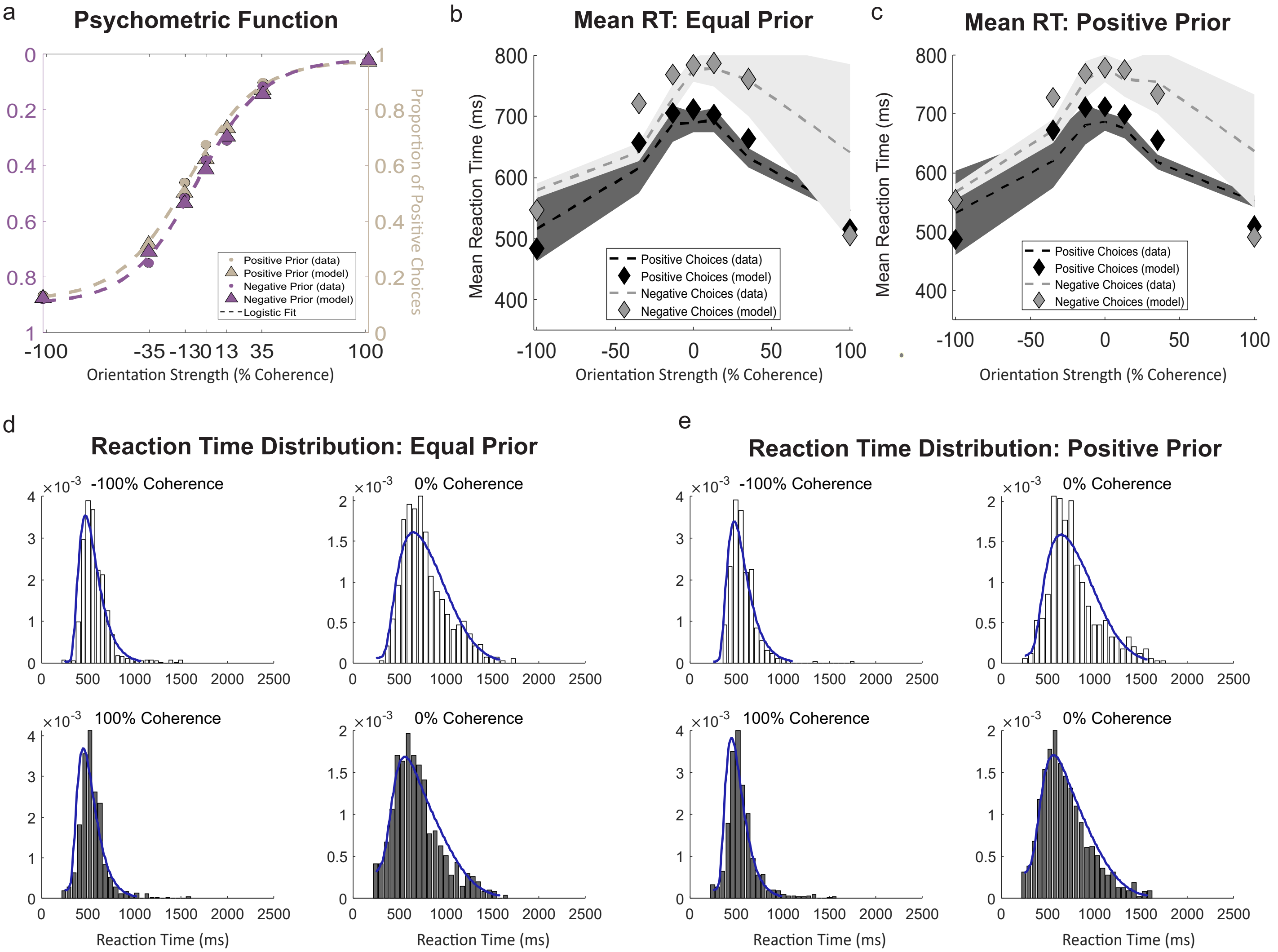


Figure S3. Model fits for choices and reaction times for Explicit condition in Experiment 1.

a. Psychometric function showing proportion of positive choices as a function of coherence for Equal prior condition (*lilac*) and Positive prior condition (*khaki*). Circles shows the data and the triangles show the model predictions and the dashed line shows the maximum likelihood logistic fit to the data.

b. Mean reaction time plot as a function of stimulus coherence for Equal prior condition. The gray color shows the mean RT for negative choices and dark color shows mean RT for positive choices. Dashed line is the mean reaction time and the shaded area shows the 95% confidence interval. The model prediction of mean RT is shown by the diamonds.

c. Same as b. for Positive Prior condition.

d. Reaction Time distributions under Equal prior condition for i) 100% coherence in negative direction and negative choices (gray, upper left), ii) 0% coherence and negative choices (gray, upper right), iii) 100% coherence in positive direction and positive choices (dark, lower left), and iv) 0% coherence and positive choices (dark, lower right). The histogram shows the data and blue traces show the model prediction.

e. Same as d. for Positive prior condition.

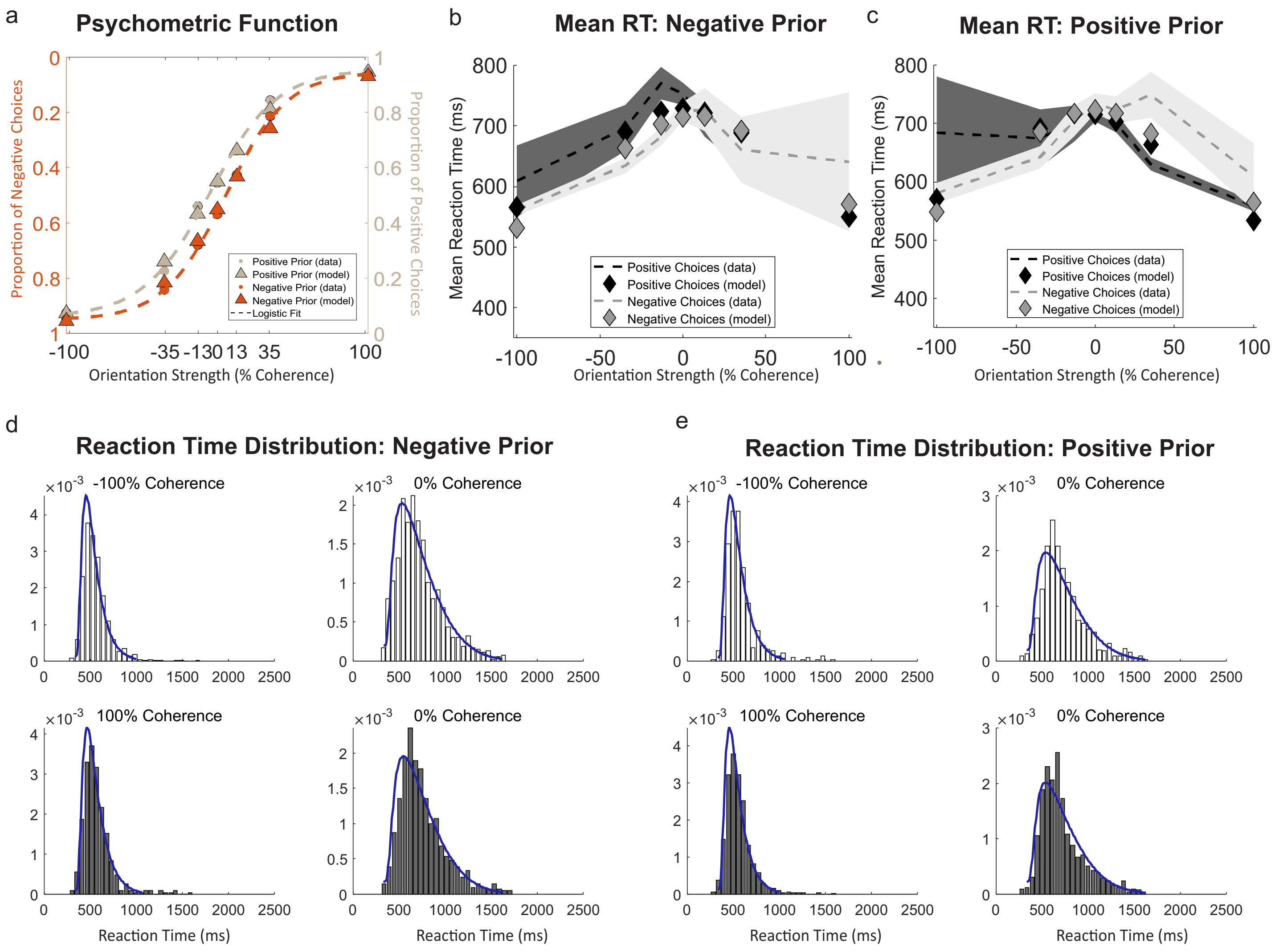


Figure S4. Model fits for choices and reaction times for Implicit condition in Experiment 2.
a. Psychometric function showing proportion of positive choices as a function of coherence for Negative prior condition (orange) and Positive prior condition (*khaki*). Circles shows the data and the triangles show the model predictions and dashed line shows the maximum likelihood logistic fit to the data.
b. Mean reaction time plot as a function of stimulus coherence for the Negative prior condition. The gray color shows the mean RT for negative choices and dark color shows mean RT for positive choices. Dashed line is the mean reaction time and the shaded area shows the 95% confidence interval. The model prediction of mean RT is shown by the diamonds.
c. Same as b. for Positive Prior condition.
d. Reaction time distributions under Negative prior condition for i) 100% coherence in negative direction and negative choices (gray, upper left), ii) 0% coherence and negative choices (gray, upper right), iii) 100% coherence in positive direction and positive choices (dark, lower left), and iv) 0% coherence and positive choices (dark, lower right). The histogram shows the data and blue traces show the model prediction.
e. Same as d. for Positive prior condition.

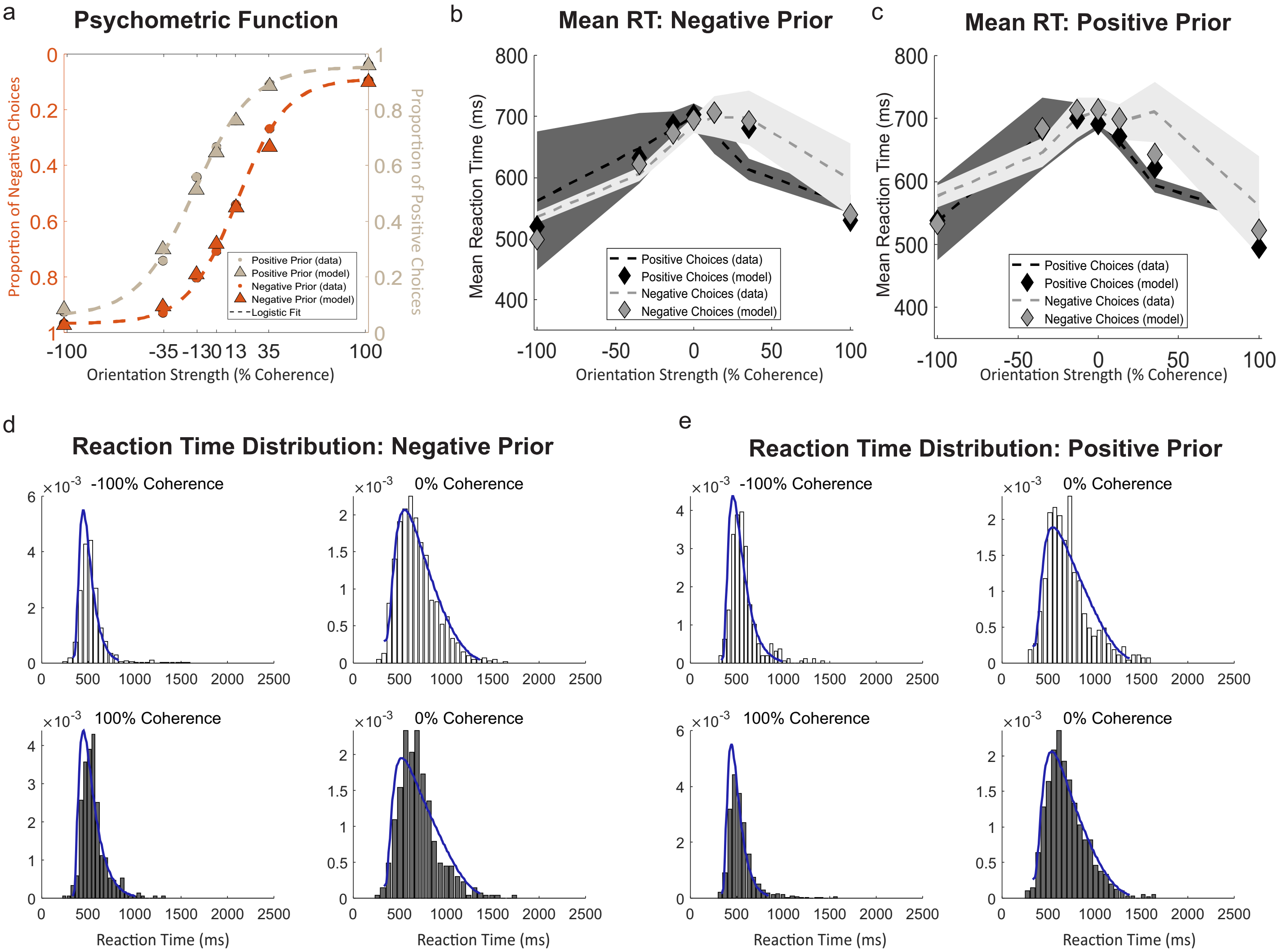


Figure S5. Model fits for choices and reaction times for Explicit condition in Experiment 2.
a. Psychometric function showing proportion of positive choices as a function of coherence for Negative prior condition (orange) and Positive prior condition (*khaki*). Circles shows the data and the triangles show the model predictions and dashed line shows the maximum likelihood logistic fit to the data.
b. Mean reaction time plot as a function of stimulus coherence for the Negative prior condition. The gray color shows the mean RT for negative choices and dark color shows mean RT for positive choices. Dashed line is the mean reaction time and the shaded area shows the 95% confidence interval. The model prediction of mean RT is shown by the diamonds.
c. Same as b. for Positive Prior condition.
d. Reaction time distributions under Negative prior condition for i) 100% coherence in negative direction and negative choices (gray, upper left), ii) 0% coherence and negative choices (gray, upper right), iii) 100% coherence in positive direction and positive choices (dark, lower left), and iv) 0% coherence and positive choices (dark, lower right). The histogram shows the data and blue traces show the model prediction.
e. Same as d. for Positive prior condition.

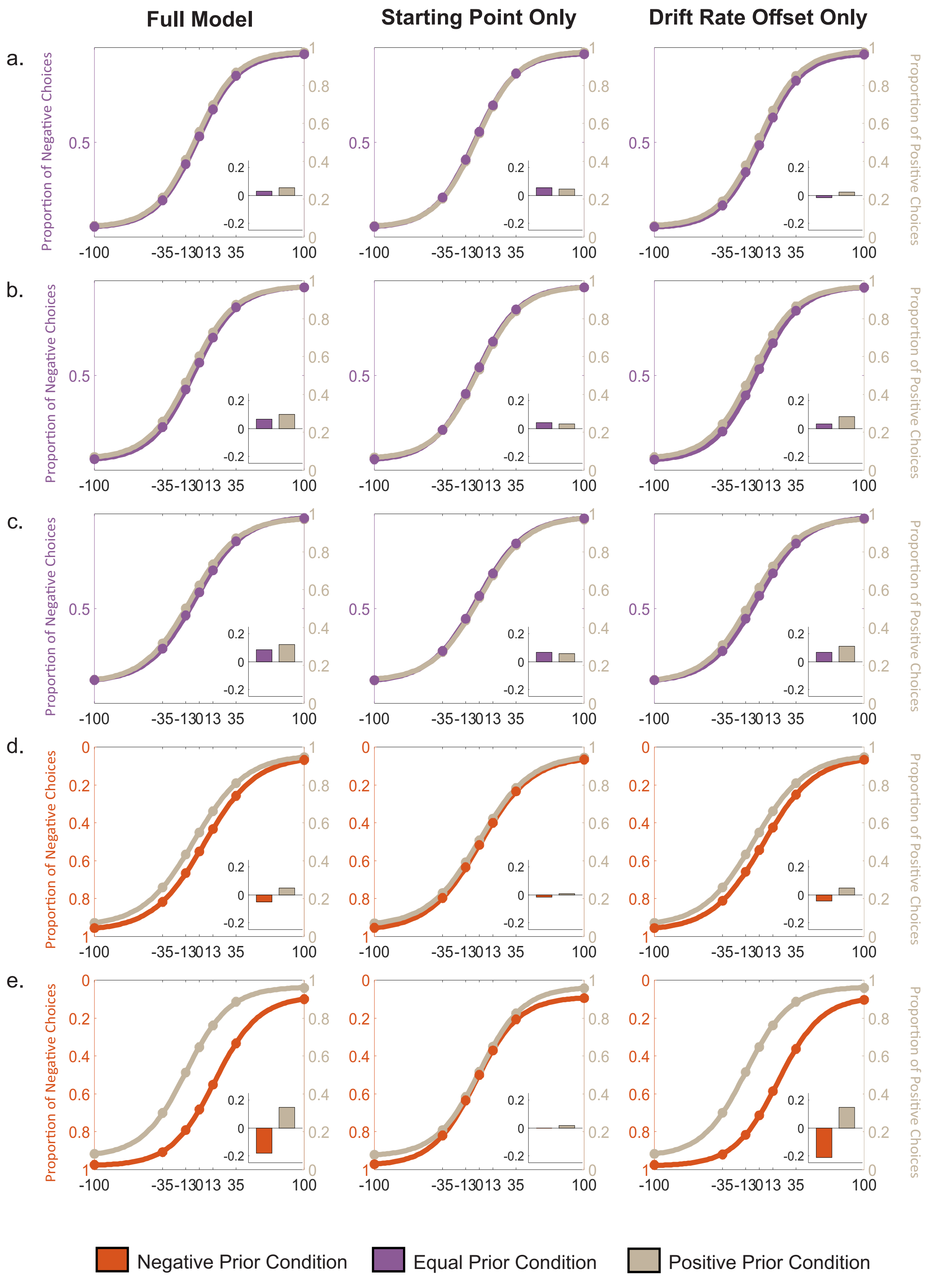


Figure S6: DDM model simulation for experiment 1 (a, b and c) and experiment 2 (d and e). a. Proportion of choices plotted against Glass pattern coherences in Implicit condition during experiment 1. The psychometric function was generated by simulating DDM model with optimized parameters including starting point offset and drift rate offset (first column), only starting point offset (second column) and only drift rate offset (third column). The inset in each figure shows the bias at 0% coherence for respective stimulus feature compared to chance level. b. same as in a for Partial Implicit condition in Experiment 1, c. same as in a for explicit condition in Experiment 1, d. same as in a for Implicit condition in Experiment 2, and e. same as in a for explicit condition in Experiment 2.

### Table S1: Model parameters for Experiment 1. Related to Figure 2.

| Parameters | | Implicit | | Partial Explicit | | Explicit | |
|---|---|---|---|---|---|---|---|
| | | *Estimate* | *S.E.* | *Estimate* | *S.E.* | *Estimate* | *S.E.* |
| **Proportionality factor between coherence and drift rate *k* (1/s·% coh) x $10^{-5}$** | | 3.3650 | 0.0442 | 4.5250 | 0.04841 | 2.525 | 0.04707 |
| **Diffusion coefficient (variance/time) ($s^{-1}$)** | | $8.8747e^{-4}$ | $3.4654e^{-5}$ | 0.0014 | $2.6232e^{-5}$ | $7.0002e^{-4}$ | $5.0839e^{-5}$ |
| **Scaling parameter for collapsing bounds *s* ($s^{-1}$)** | | 0.0047 | $9.909e^{-5}$ | 0.0030 | 2.6076 | 0.0033 | $1.8472e^{-4}$ |
| **Delay parameter for collapsing bounds *d* (ms)** | | 81.0029 | 87.3539 | 1 | 34.6458 | 241 | 34.3183 |
| **Residual time $t_0$ (ms)** | | 309 | 1.8568 | 326 | 1.9021 | 236.9997 | 2.0751 |
| **Equal Prior Condition** | **Starting point offset** | 0.0755 | 0.007 | 0.0636 | 0.0080 | 0.0318 | 0.0075 |
| | **Drift-rate offset *O* ($s^{-1}$) x $10^{-5}$** | -7.8201 | 1.7731 | 9.9751 | 2.2828 | 5.0000 | 1.5372 |
| | **Probability of positive response** | 0.0517 | 0.0059 | 0.0519 | 0.0064 | 0.1170 | 0.0050 |
| | **Probability of negative response** | 0.0294 | 0.0063 | 0.0283 | 0.0065 | 0.0155 | 0.0069 |
| **Positive Prior Condition** | **Starting point offset** | 0.0504 | 0.0070 | 0.0315 | 0.0080 | 0.0215 | 0.0076 |
| | **Drift-rate offset *O* ($s^{-1}$) x $10^{-5}$** | 2.499 | 1.7689 | 29.698 | 2.3150 | 17.50 | 1.5122 |
| | **Probability of positive response** | 0.0559 | 0.0058 | 0.0629 | 0.0063 | 0.1166 | 0.0052 |
| | **Probability of negative response** | 0.0202 | 0.0066 | 0.0284 | 0.0063 | 0.0218 | 0.0064 |
| **Mean of RT distribution for uninformed decision** | | 51.3709 | 18.421 | 26.3029 | 24.977 | 194.1021 | 10.9023 |
| **SD of RT distribution for uninformed decisions** | | 196.6445 | 8.6075 | 236.7432 | 10.8925 | 172.3235 | 4.8731 |

**Table S2: Model parameters for Experiment 2. Related to Figure 3.**

| Parameters | | Implicit | | Explicit | |
|---|---|---|---|---|---|
| | | *Estimate* | *S.E.* | *Estimate* | *S.E.* |
| **Proportionality factor between coherence and drift rate *k* (1/s·% coh) x $10^{-5}$** | | 3.7125 | 0.0498 | 4.40 | 0.049153 |
| **Diffusion coefficient (variance/time) ($s^{-1}$)** | | 0.0014 | $2.837e^{-5}$ | 0.0014 | $2.6619e^{-5}$ |
| **Scaling parameter for collapsing bounds *s* ($s^{-1}$)** | | 0.0026 | $3.2151e^{-4}$ | 0.0027 | $3.7582e^{-4}$ |
| **Delay parameter for collapsing bounds *d* (ms)** | | 1 | 38.576 | 480.996 | 35.8201 |
| **Residual time $t_0$ (ms)** | | 316 | 2.0365 | 312 | 1.877 |
| **Negative Prior Condition** | **Starting point offset** | -0.0159 | 0.0081 | 0.0686 | 0.0084 |
| | **Drift-rate offset *O* ($s^{-1}$) x $10^{-5}$** | -13.781 | 2.1591 | -78.750 | 2.4168 |
| | **Probability of positive response** | 0.0367 | 0.0065 | 0.0238 | 0.0062 |
| | **Probability of negative response** | 0.0510 | 0.0067 | 0.0883 | 0.0062 |
| **Positive Prior Condition** | **Starting point offset** | $0.0001e^{-3}$ | 0.008 | -0.0031 | 0.0084 |
| | **Drift-rate offset *O* ($s^{-1}$) x $10^{-5}$** | 17.500 | 2.2086 | 55.781 | 2.3808 |
| | **Probability of positive response** | 0.0600 | 0.0066 | 0.0754 | 0.0065 |
| | **Probability of negative response** | 0.0419 | 0.0064 | 0.0354 | 0.0059 |
| **Mean of RT distribution for uninformed decision** | | 128.0017 | 25.3396 | 135.2723 | 17.7876 |
| **SD of RT distribution for uninformed decisions** | | 247.4817 | 8.5055 | 200.3777 | 7.4011 |

**Table S3: Model comparisons between model with only start point offset, model with only drift-rate offset, and model with both starting point and drift-rate offset. Related to Figure 2 and 3**

| BIC scores | | Only starting point offsets | Only drift-rate offsets | Both types of offsets |
|---|---|---|---|---|
| **75-50 Experiment** | **Implicit** | 67193 | 67255 | 66982 |
| | **Partial Explicit** | 68162 | 68034 | 67892 |
| | **Explicit** | 71179 | 71056 | 70695 |
| **75-25 Experiment** | **Implicit** | 70915 | 70898 | 70679 |
| | **Explicit** | 68296 | 68084 | 68136 |
